# The structure-fitness landscape of pairwise relations in generative sequence models

**DOI:** 10.1101/2020.11.29.402875

**Authors:** Dylan Marshall, Haobo Wang, Michael Stiffler, Justas Dauparas, Peter Koo, Sergey Ovchinnikov

## Abstract

If disentangled properly, patterns distilled from evolutionarily related sequences of a given protein family can inform their traits - such as their structure and function. Recent years have seen an increase in the complexity of generative models towards capturing these patterns; from sitewise to pairwise to deep and variational. In this study we evaluate the degree of structure and fitness patterns learned by a suite of progressively complex models. We introduce pairwise saliency, a novel method for evaluating the degree of captured structural information. We also quantify the fitness information learned by these models by using them to predict the fitness of mutant sequences and then correlate these predictions against their measured fitness values. We observe that models that inform structure do not necessarily inform fitness and vice versa, contrasting recent claims in this field. Our work highlights a dearth of consistency across fitness assays as well as divergently provides a general approach for understanding the pairwise decomposable relations learned by a given generative sequence model.

## 1 Introduction

Inferring biophysical characteristics of a biological sequence from sequence alone is an outstanding challenge in computational biology. By comparing homologous sequences, patterns associated with such characteristics can emerge - thus allowing one to infer them if a sufficiently representative set of sequences are available. Non-exhaustively, these patterns include: *conservation* - per-position frequencies indicative of function; *coevolution* - covariation between positions often in physical residue-residue contact; and *phylogeny* - relationship between sequences akin to the organization of species from which the sequences were sourced. See Fig. 1A. Among other reasons, disentanglement of the origins of these patterns is desirable because better resolved coevolution and phylogeny could improve structure prediction and species delimitation, respectively. It is challenged, however, by the fact that they confound one another and is an unsolved problem of the field.

**Figure 1:**
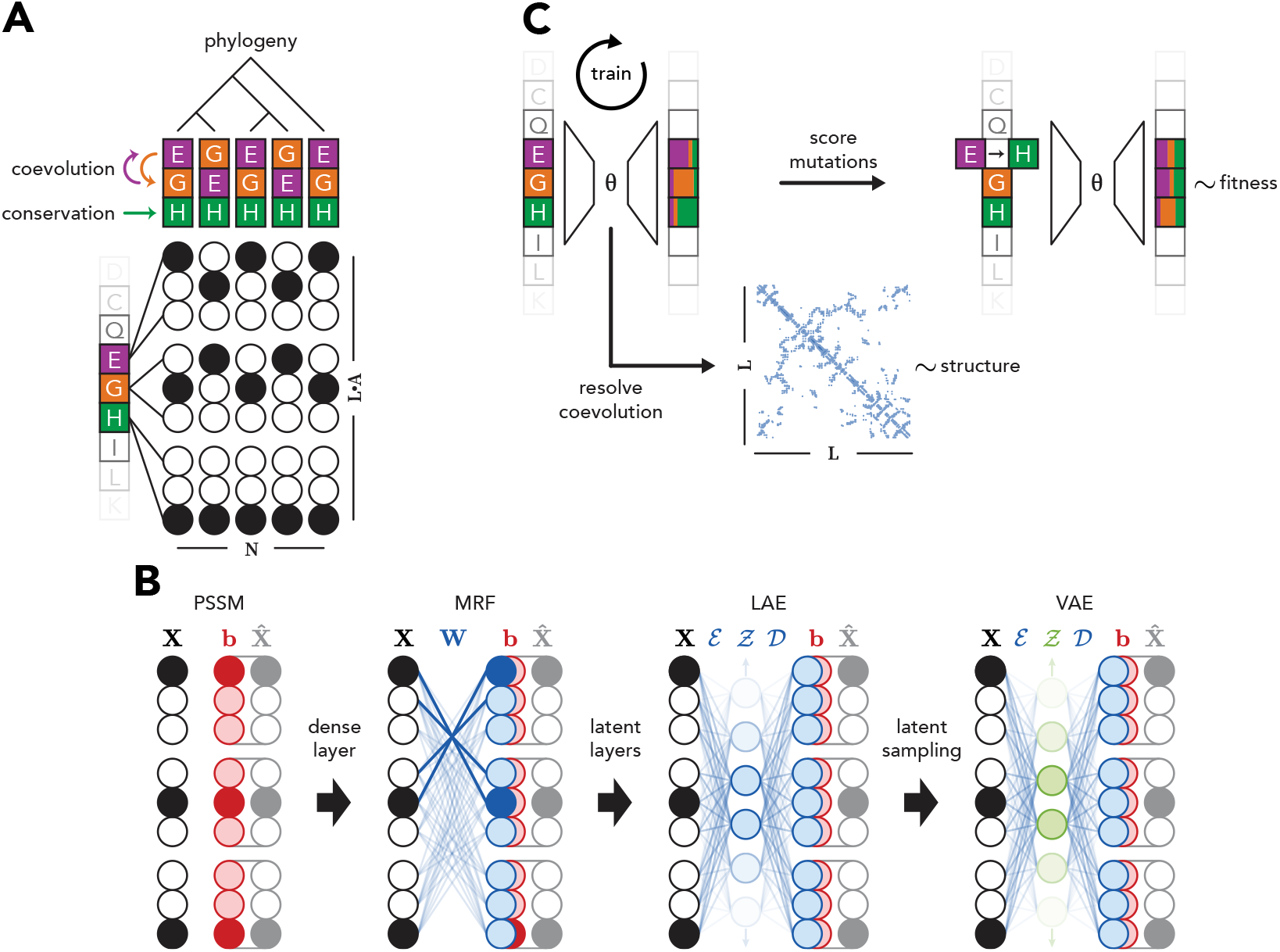
Patterns within and between homologous sequences can inform the structure and fitness of a representative protein therein. **(A)** Schematic of a one-hot encoded MSA in matrix form. Patterns of interest: conservation, coevolution, and phylogeny. **(B)** Iterative complexity of considered generative sequence models. **(C)** Prediction tasks of trained models: fitness ~ mutation effect; structure residues in contact.

Early generative sequence models, such as Position-Specific Scoring Matrices (PSSMs) [Stormo et al., 1982], were able to distinguish some functional characteristics but were limited to only considering sitewise relations. Later, Markov Random Fields (MRFs) were applied to resolve molecular coevolution by incorporating pairwise relations [Lapedes et al., 1999, Thomas et al., 2008, Weigt et al., 2009, Balakrishnan et al., 2011, Morcos et al., 2011, Jones et al., 2012, de Juan et al., 2013]. Improvements, such as pseudolikelihood maximization [Ekeberg et al., 2013, Kamisetty et al., 2013], have since elevated MRFs, resulting in dramatically improved structure prediction. Indeed, in addition to other inputs, MRF features underpin AlphaFold, the reigning Critical Assessment of protein Structure Prediction (CASP) champion [Senior et al., 2020].

Predicting the functional effect of mutant variants for a given protein by unsupervised means has seen renewed interest in recent years. The strategy entails using the aforementioned models to score mutant sequences relative to a wildtype sequence and then compare the scores against measured phenotypes of these mutants. An approach built on a coevolution model was initially proposed by Lapedes et al. [2002] to predict the thermostability of Fyn SH3 domain mutants - as quantified by ΔΔ*G_mutant_*. This idea was again demonstrated later in Figliuzzi et al. [2016] but for predicting the fitness effect of mutant TEM-1 Beta Lactamase sequences instead - as quantified by a specific enzymatic selection assay. Hopf et al. [2017] built on this, generalizing the method to more proteins. Riesselman et al. [2018] claimed state of the art performance for this task with a deep Variational Autoencoder (VAE). They also claimed that by dint of its improved performance over pairwise models that it necessarily captured higher-order dependencies. It is, however, unclear if the model is actually learning higher-order interactions or simply a mixture of PSSMs where differentially conserved functional positions - between different groups of sequences - would be expected to cluster together in the structure.

Contemporary to this, other work also applied VAEs to fitness inference. Notably, both Sinai et al. [2017] and Ding et al. [2019] allude to their VAEs capturing varying levels of conservation - within groups of phylogenetically related sequences - as opposed to coevolution. Arising from this ambiguity came a natural question: what exactly are these models learning that results in their differing ability to infer function and structure? In parallel, what role do pairwise relations play in this task?

In this work, we attempt to address these questions. We focus on the widely studied TEM-1 Beta Lactamase, the central case study of Figliuzzi et al. [2016], Hopf et al. [2017], and Riesselman et al. [2018]. We carefully resolve how structure and mutant fitness are related on an experimental level. We systematically evaluate a range of generative sequence models from PSSM to VAE by using them to predict these two properties for TEM-1 Beta Lactamase. We assess a wide range of model hyperparameters. We take stock of how prediction and experiment relate to one another. We observe that models perform these tasks differentially rather than commensurately. We conclude that patterns in homologous sequences that inform structure differ from those that inform fitness. Paper code: https://github.com/sokrypton/seqsal_v2.

## 2 Data

Here we describe the considered sequence, structure, and fitness data of TEM-1 Beta Lactamase, the case study protein. For brevity, ***β_ℓ_***:= TEM-1 Beta Lactamase. ***β_£_*** is an enzyme capable of hydrolyzing penicillin type β-lactam antibiotics [Abraham and Chain, 1940]. It exhibits desirable features that confer greater confidence in inferring biophysical characteristics solely from homologous sequence comparison. It is monomeric, globular, purportedly singular in function, has no known cofactors, and is found primarily on plasmids [Stiffler et al., 2015, Naas et al., 2017, Bush, 2018]. Thus, disentanglement of the structure-fitness landscape from sequence alone in this protein is less confounded than in most other proteins.

### 2.1 Sequence

#### 1 Training data

The ***β_ℓ_*** sequences were organized in a multiple sequence alignment (MSA); a set of *N* homologous sequences with alphabet *A* and length *L*. In one-hot encoded form: MSA:= **X** ∈ {0, 1}^*N*×*L*×*A*^, as shown in Fig. 1A. The set *A* represents the 20 amino acids as well as a category for gaps *g*. We sourced the MSA from Riesselman et al. [2018], which was generated using jackhmmer [Eddy, 2011] and pulled sequences from UniRef100 [Suzek et al., 2015]. The reference (or query) sequence of interest **r**_*L*×*A*_ ∈ *X* is the wildtype ***β_ℓ_*** sequence.

#### Testing data

Starting from the same reference sequence, **r**_*L×A*_ ∈ *X*, we also generated another MSA **Y** ∈ {0, 1}^*N*^_*Y*_ ^×*L*×*A*^. `HMMER` [Eddy, 2011] with a bit score of 27 was used to pull sequences from a metagenomics database as described in Ovchinnikov et al. [2017].

#### Both datasets

For both MSAs, sequences ≥ 20% gapped positions relative to the query sequence and that shared ≥ 90% sequence identity to each other were removed. Additionally, sequences in *Y* sharing ≥ 90% sequence identity with any sequence in *X* were also removed.

### 2.2 Structure

A set of ***β_ℓ_*** labeled x-ray crystal structures was sourced from the Beta-Lactamase DataBase (BLDB) [Naas et al., 2017], a manually curated collection in turn sourced from the Protein Data Bank (PDB) [Berman et al., 2000, Burley et al., 2019]. Structures were not considered further if the correspondent sequence could not be one-hot encoded given the aforementioned alphabet *A*, such as those with noncanonical amino acids. The resulting forty-two ***β_ℓ_*** structures were processed with ConFind [Zheng et al., 2015] to identify physically interacting residues. Each structure is represented by a contact map,

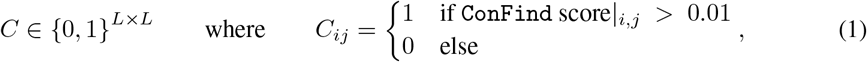

as recommended by Zheng et al. [2015]. See Fig. 1C.

### 2.3 Fitness

Data that assessed varying aspects of ***β_ℓ_*** fitness and function were collected. The nine datasets were generated by experimentally or computationally characterizing single mutant variants of the ***β_ℓ_*** reference sequence **r**_*L*×*A*_. These mutant sequences 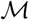 are defined as all possible missense mutations *m* for each sequence index *i* ∈ {1,…, *L*}; and more formally in one-hot encoded form,

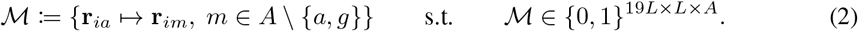

The fitness assay datasets, denoted by 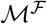, are a mapping from the mutant sequences. They represent the observed phenotypes for a given fitness assay 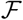 over some subset of 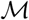,

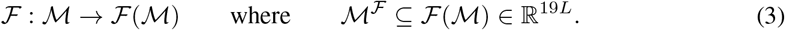

## 3 Models

### 3.1 Generative sequence models

Inspired by Dauparas et al. [2019], the investigated models progress in complexity from trivial to highly non-convex. A graphical schematic is shown in Fig. 1B. Each learn a different composition of biological relations within a MSA. All models *f* learn to reconstruct a MSA 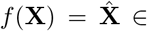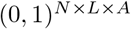 with categorical cross entropy

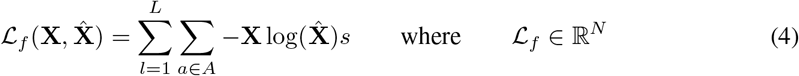

as the minimized loss function. In the following, softmax is taken along the alphabet *A* axis and model weights were *L*_2_ regularized with coefficient *λ*.

#### 3.1.1 Position-Specific Scoring Matrix (PSSM)

PSSMs capture the sitewise decomposable relations within **X** representing evolutionary conservation [Stormo et al., 1982]. They are parameterized by a bias matrix **b** ∈ ℝ^*L*×*A*^. As noted in Dauparas et al. [2019], given

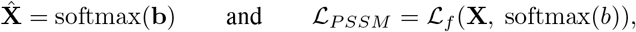

the analytically derived solution for **b** is 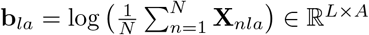.

#### 3.1.2 Markov Random Field (MRF)

MRFs with the pseudolikelihood approximation [Balakrishnan et al., 2011, Ekeberg et al., 2013, Kamisetty et al., 2013] capture the patterns within **X** representing coevolution [Lapedes et al., 1999, Weigt et al., 2009, Morcos et al., 2011]. They are parameterized by an explicitly pairwise decomposable weight matrix, **W** ∈ ℝ^*L*×*A*×*L*×*A*^, and a bias matrix, **b**∈ ℝ^1×*L*×*A*^. The trivial residue self-mapping is precluded by the constraint **W**_*i*:*i*:_ → 0 | *i* ∈ {1,…,*L*}. Given

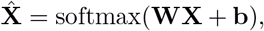

the MRF loss function 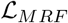 is *L*_2_ regularized with coefficient *λ*

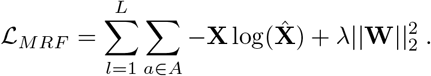

#### 3.1.3 Linear Autoencoder (LAE)

LAEs [Baldi and Hornik, 1989, Kunin et al., 2019] are a flexible model type capable of capturing varying relations within **X**. In this work, they share the same framework as MRFs but differ through the inclusion of latent linear layers *l*. The architecture is s.t. **X** is first encoded to latent space 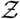, where 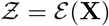, with encoder 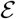. Next, it is decoded to 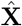, where

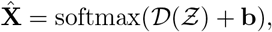

with decoder 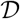. Whereas the pairwise relations are explicitly parameterized in the MRF, they are unknown for the LAE. The LAE loss function 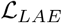 is defined as

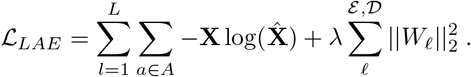

#### 3.1.4 Variational Autoencoder (VAE)

Similar to LAEs, VAEs [Kingma and Welling, 2013] are a flexible model type. In this work, they share the same framework as LAEs but differ by: using selu [Klambauer et al., 2017] non-linear activation, `dropout` [Srivastava et al., 2014], and `batch normalization` [Ioffe and Szegedy, 2015] for 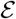 and 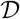, and latent probabilistic sampling

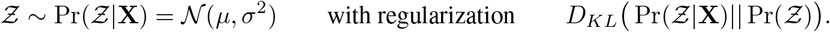

Riesselman et al. [2018] claim the capture of triwise (or higher) relations with a VAE but do not assess its learned pairwise relations which remain unresolved. Given

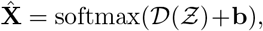

the VAE loss function 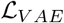 is defined as

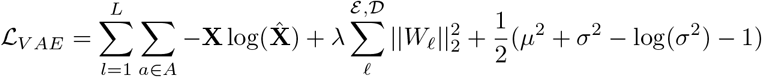

where the Kullback-Leibler divergence regularization term is written in the alternative estimator form, as derived in Kingma and Welling [2013].

### 3.2 Training approach

Models were trained to reconstruct **X** over a range of combinatorily sampled hyperparameters. The possible hyperparameters increase from PSSM to MRF to LAE to VAE. The PSSM was analytically solved. MRFs, which are convex, converged to a global minima. The non-convex LAEs and VAEs were trained with batch and epoch scheduling. Where applicable, *L*_2_ regularization coefficient *λ* was uniformly sampled over (0, 1); the number of possible logits in 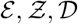 spanned base 2 from 2^8^ to 2^12^; dropout was uniformly sampled over [0, 0.5]; batch sizes were scheduled in base 2 from 2^6^ to 2^12^; and number of epochs for each batch size increment was randomly chosen over a base 2 range from small 2^3^ to large 2^6^. Parameters were optimized with the default `Keras` [Chollet et al., 2015] Adam optimizer [Kingma and Ba, 2014].

## 4 Pairwise saliency

For a given generative sequence model *f*, pairwise saliency **P** quantifies the composition of learned pairwise decomposable relations. We define **P** as the symmetrized Jacobian **J** evaluated with input **0**_1×*L*×*A*_. Symmetry is achieved by averaging the Jacobian and its transpose

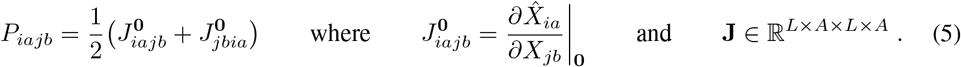

This method is conceptually similar to linearization of a non-linear model by the first order Taylor approximation evaluated around the origin.

Resolving inter-residue contacts from pairwise decomposable relations is accomplished by a variant of Average Product Correction (APC) of **P**[Dunn et al., 2008], which filters out low-rank artifacts. The diagonal of **P** before and after APC is set to 0. A visualization of the first step of APC, where the *L*_2_ norm of **P** is taken over the alphabet axes *A* not including gaps

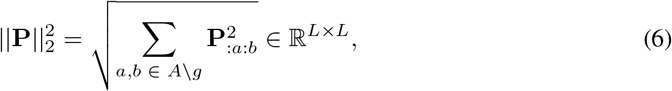

is shown for selected MRF, LAE and VAE models in Fig. 2 (**top**). The APC of **P** is shown in Fig. 2 (**middle**). An overlay of the pairwise saliency derived contacts *Ĉ* on ground truth contacts *C* is shown in Fig. 2 (**bottom**).

## 5 Data

### 5.1 Metrics between measurements and predictions

#### Quantifying structure

Contact AUC quantifies how well an alternative or predicted contact map *Ĉ* matches a ground truth contact map *C*. It is the average of the precision PPV evaluated over the highest ranking predicted contacts Ĉ_*ij*_, constrained to |*i* − *j*| > 6, in *L/*10 increments,

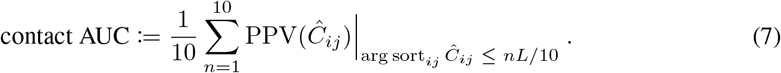

**Figure 2:**
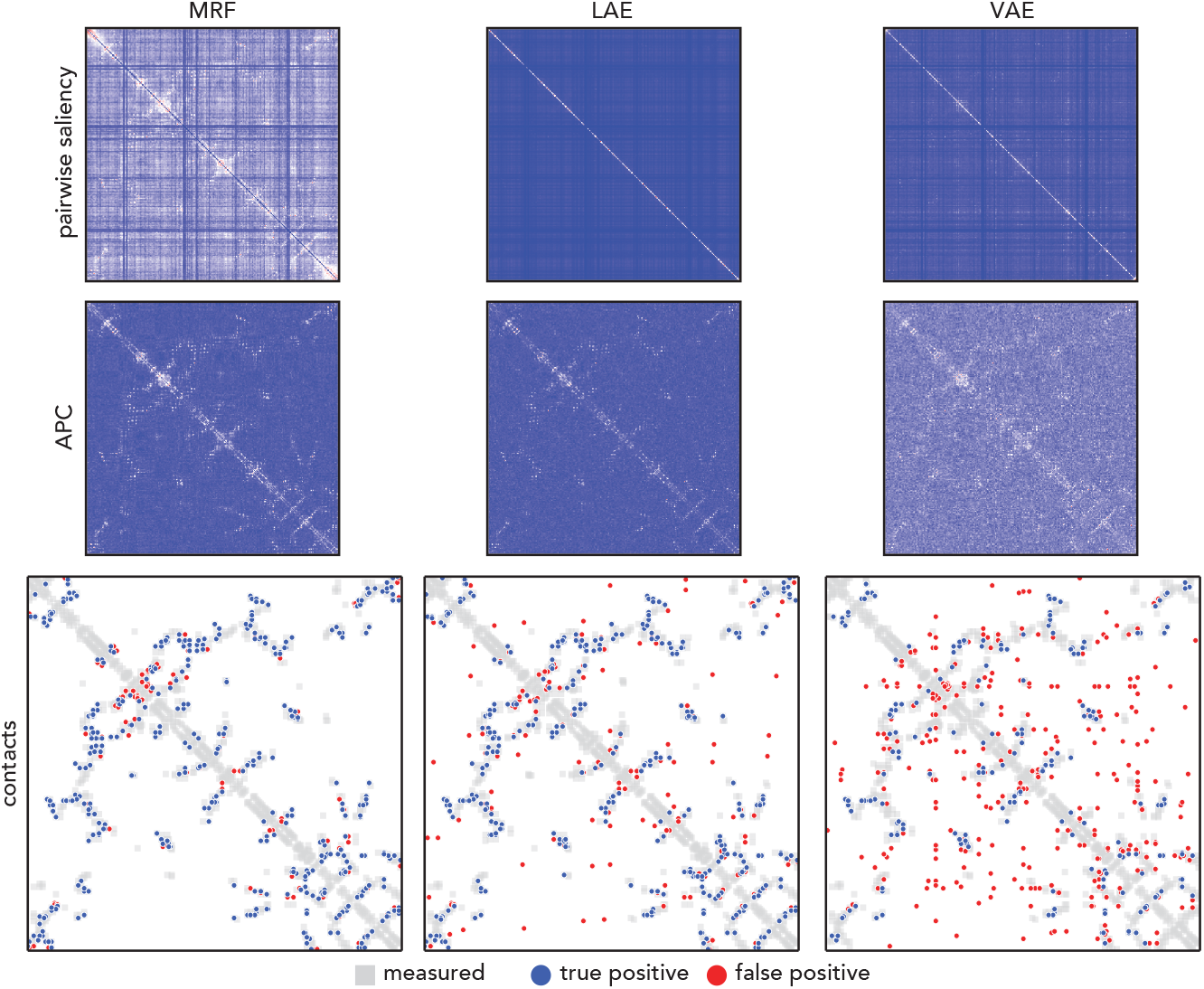
Pairwise saliency reveals structurally relevant pairwise decomposable relations learned by a given generative sequence model. **(top)***L*_2_ norm of pairwise saliency matrix over alphabet axes. **(middle)** Average product corrected (APC) pairwise saliency matrix. **(bottom)** Top *L* contacts at least 6 sequence indices apart derived from APC pairwise saliency matrix overlayed on ground truth contacts (PDB ID: 1ERO [Ness et al., 2000]).

The degree of structural information learned for a given model *f* considers the APC form of the pairwise saliency matrix **P**.

#### Quantifying fitness

Fitness has many definitions across many fields. In this work, fitness is contextually defined as the mapping from each possible missense mutation per sequence index to an assay-specific scalar, 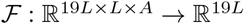, as described in Eqns 2 and 3.

First, a trained model *f* reconstructs the mutant sequences 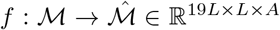. Next, the reconstruction cost is calculated with 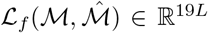, as defined by Eqn. 4. These are the predicted mutation effect values. Note that from an unsupervised model perspective, there can only be one prediction per mutation because reconstruction cost is a function of 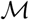 and not 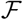. Subsequently, predicted mutation effects are correlated against 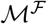, the fitness assay datasets, via absolute value of the rank (Spearman) correlation

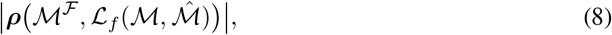

for a selected 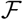, building on the precedent set by Hopf et al. [2017] and Riesselman et al. [2018]. See Fig. 3A.

**Figure 3:**
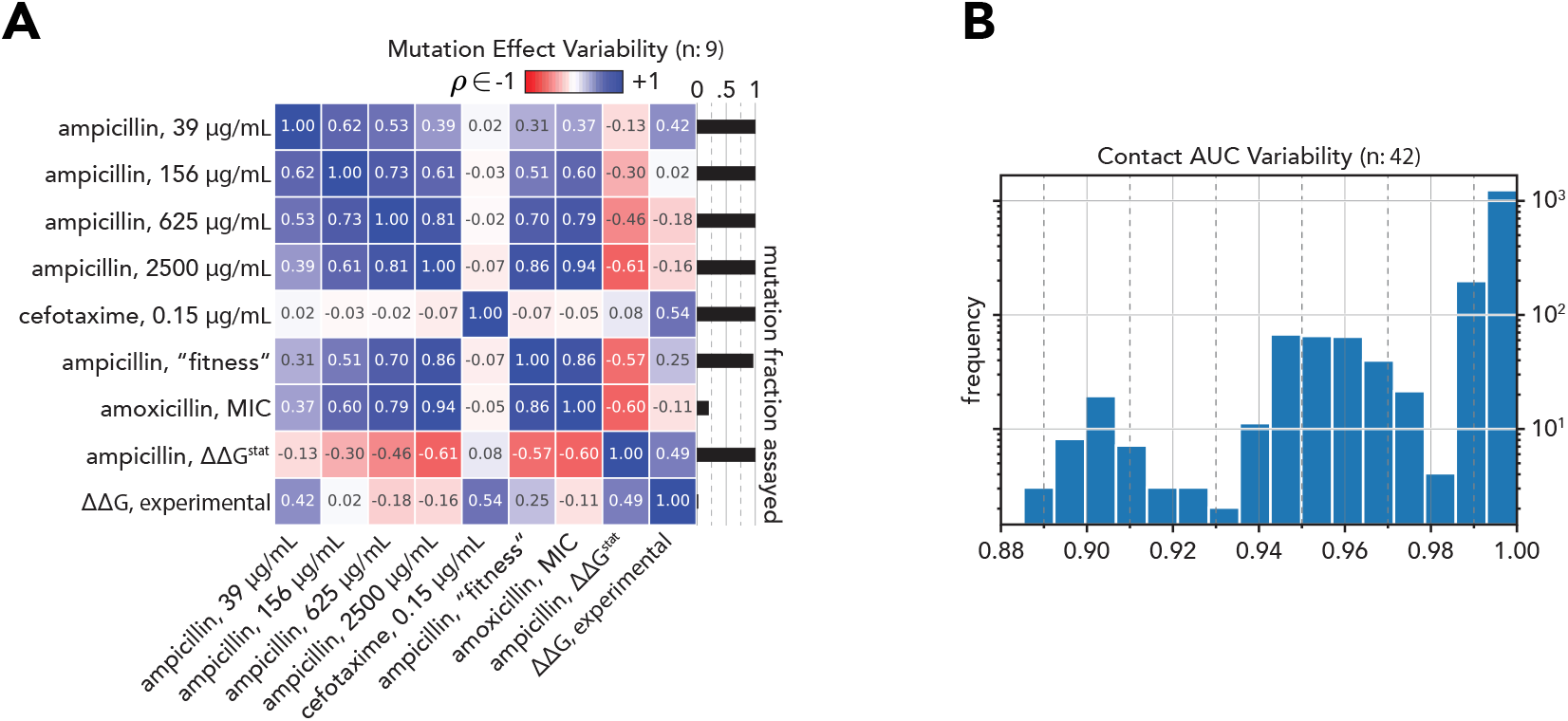
“Ground truth” varies for mutation effect but only slightly for structural contacts in TEM-1 Beta Lactamase. **(A)** All by all rank correlations [1, 1] between 9 mutation effect assays. **(B)** All by all contact AUC values [0, 1] between 42 PDB structures curated by the Beta-Lactamase DataBase (BLDB).

### 5.2 Meaning and context

#### Structure is an invariant biophysical characteristic

Contact AUC between each of the 42 ***β_ℓ_*** PDB structures varies minimally, with span ∈ [0.885, 1.000]. See Fig. 3B. This suggests quantification of contacts by contact AUC is not subjective for this protein.

#### Mutation effect informs fitness relatively

Nine mutation effect datasets interrogating function and thermostability were compared. Considering ***β_ℓ_*** is a penicillin hydrolyzing enzyme, any of "ampicillin, [39, 156, 625, 2500] μg/mL” [Stiffler et al., 2015], “ampicillin, fitness” [Firnberg et al., 2014], “amoxicillin, MIC” [Jacquier et al., 2013], “ampicillin, ΔΔG^stat^ “ [Deng et al., 2012], can claim to be representative of mutation effect when the only consideration is how such mutations affects ***β_ℓ_***’s ability to hydrolyze its natural substrate. While “ampicillin, 2500 μg/mL” [Stiffler et al., 2015] has been the ground truth assay of choice for previous work using unsupervised models to infer mutation effect for ***β_ℓ_*** [Hopf et al., 2017, Riesselman et al., 2018], and correlates strongly with “ampicillin, fitness” [Firnberg et al., 2014] and “amoxicillin, MIC” [Jacquier et al., 2013], it negatively correlates with “ampicillin, ΔΔG^stat^” [Deng et al., 2012], and variably correlates with the same assay from the same paper but at lower concentrations “ampicillin [39, 156, 625] μg/mL” [Stiffler et al., 2015]. Similarly, if the consideration is instead how a mutation impacts ***β_ℓ_***’s ability to hydrolyze “cefotaxime, 0.15 μg/mL”, a different *β*-lactam antibiotic [Stiffler et al., 2015], or thermostability [Yang et al., 2020] - the outcomes also vary.

Despite being singular in purpose, to break down a single class of molecules that inhibit bacterial cell wall formation, ***β_ℓ_*** mutation effect varies under selection of its natural substrate. It also depends on the consistency of method for calculating mutation effect from raw allele count ratios. While this subjectivity is alluded to in previous work [Lapedes et al., 2002, Figliuzzi et al., 2016, Hopf et al., 2017, Riesselman et al., 2018], it was not explored in depth. It is possible there are better metrics than rank correlation for this task. We note contemporary efforts to standardize the terminology and metrics for this data type [Esposito et al., 2019, Dunham and Beltrao, 2020], but consensus has yet to emerge.

## 6 Results

Taking scope of the evaluated generative sequence models over a range of hyperparameters, a view of the structure-fitness landscape comes into focus. Predicted structure *Ĉ* is resolved through pairwise saliency **P** (Eqn 5) and compared to ground truth by contact AUC (Eqn 7). Structural ground truth is PDB ID 1ERO [Ness et al., 2000]. Note that 1ERO can be substituted by any of the other structures in Fig. 3B. Predicted mutant effect function 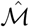 is determined by |***ρ***| (Eqn 8). “Ampicillin, 2500 μg/mL” [Stiffler et al., 2015] represents ground truth for mutant effect fitness, chosen as a consequence of precedent [Hopf et al., 2017, Riesselman et al., 2018] and also because it reasonably represents ***β_ℓ_***’s natural function. Generalizability is proxied by the reconstruction cost 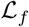 (Eqn 4) of the predicted test data Ŷ, revealing how underfit or overfit *f* is on **X**.

Shown in Fig. 4A is the relation between task performance for the generative sequence models across the aforementioned hyperparameters. Each circle is an individual model; ensembling is not considered. By definition a PSSM, purple circle, does not capture pairwise relations. MRFs, red circles, infer structure to almost within experimental error and infer fitness decently. LAEs, green circles, span the spectrum of both tasks, but not as well as MRFs for structure or VAEs for fitness. VAEs, blue circles, learn fitness well and learn structure variably. Also plotted is DeepSequence [Riesselman et al., 2018] for comparison, orange circle, reproduced from weights provided online. Shown in Fig. 4B is the relation between learned structure and test loss. Note that arg max_*f*_ (contact AUC) = arg max_*f*_ (test loss) for both LAE and VAE.

**Figure 4:**
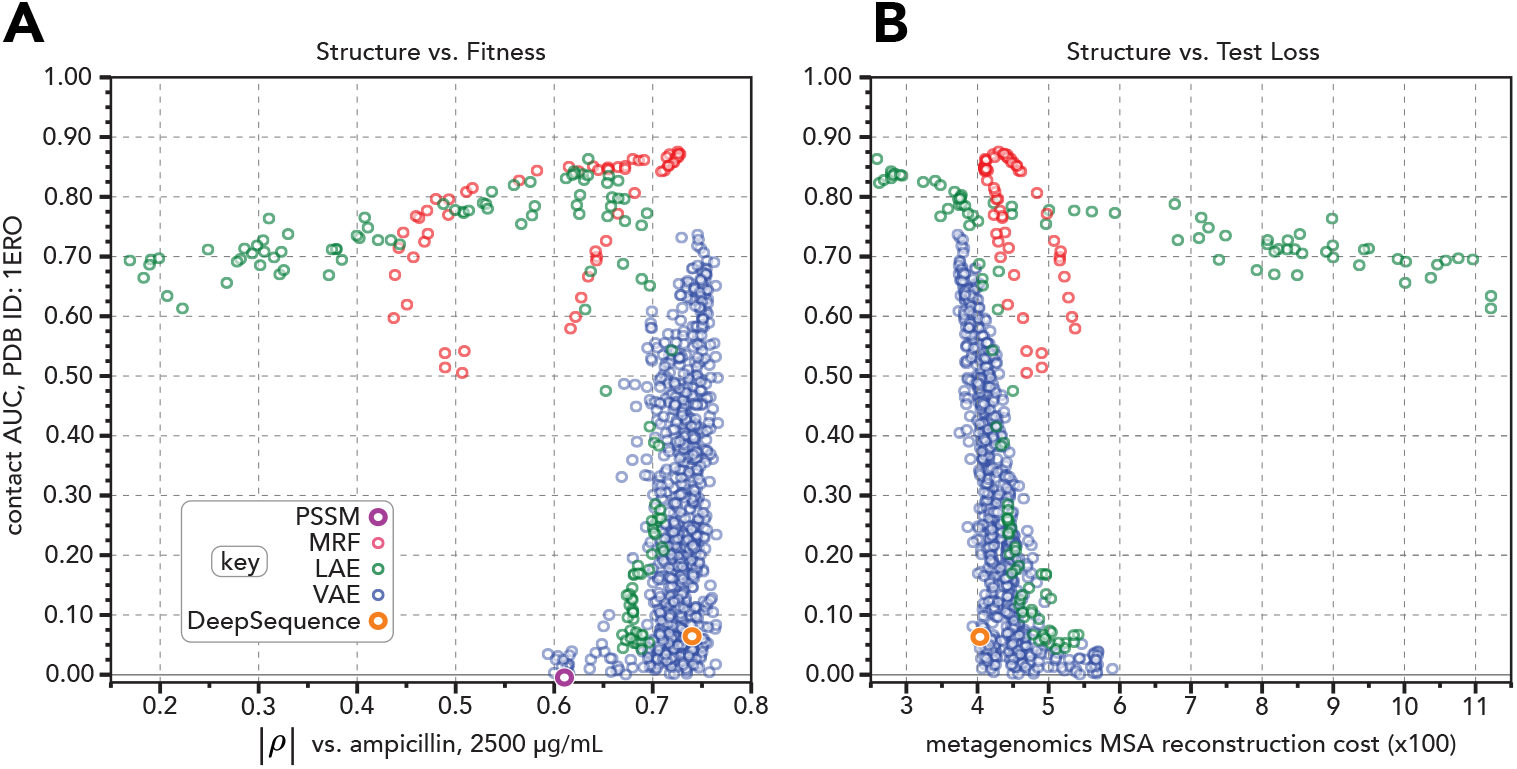
Generative models learn structure and fitness differentially. High throughput evaluation of model performance towards inferring structure (y-axis) versus **(A)** inferring mutation effect for a representative fitness assay and **(B)** reconstructing a metagenomics derived homologous MSA.

We therefore conclude that models can infer structure and fitness differentially. We also conclude that within each model category, models that best reconstruct the test data tend to learn contacts the best.

## 7 Discussion

In this investigation on the structure-fitness landscape of TEM-1 Beta Lactamase, we clarify the relationship between model parameterization and captured biological information for a suite of progressively complex generative sequence models. We introduce a novel method, pairwise saliency, to reveal the degree of structure they have learned. We also assess their capacity to infer fitness, proxied by the measured effect of mutant sequences. Surprisingly, we find that models can learn one task and not the other. It is possible pairwise saliency insufficiently resolves learned contacts for complex models. It is also possible the mutant effect data does not represent the aggregate evolutionary pressures etched into the patterns found across homologous ***β_ℓ_*** sequences. We are also left wondering how relevant intra-sequence dependencies are for fitness inference in an unsupervised framework.

We suspect that models that infer mutation effect well but structure poorly are learning mixture models, where each group of sequences emit a single sequence profile. From our analyses, it seems possible to create a hybrid model that learns both coevolution along with a hierarchical mixture bias term for phylogeny. Such a model could both better predict structure as well as more accurately delimit clades within the tree of life. It is also unclear what, exactly, higher-order relations are - as previous work claims to have captured [Riesselman et al., 2018]. Is it possible that phylogeny itself are the higher-order relations in question? Previous work has shown that active sites within proteins can be predicted by scoring how well each position of a multiple sequence alignment agrees with the overall phylogenetic gene tree [La et al., 2005].

Model interpretability continues to be a heavily debated topic that lacks consensus [Gilpin et al., 2018]. This in mind we simply propose pairwise saliency merely as a starting point for further study into the disentangled relations resolved from generative sequence models. Indeed, immediately pursued next steps include but are not limited to: factorizing the sitewise terms that are theoretically confounding pairwise saliency, utilizing the Hessian towards distinguishing triwise decomposable relations, and perhaps even applying it to large natural language processing based protein sequence models that are currently in vogue [Rao et al., 2019, Alley et al., 2019, Rives et al., 2019, Elnaggar et al., 2020].

## Acknowledgments and Disclosure of Funding

SO is supported by the John Harvard Distinguished Science Fellows Program within the FAS Division of Science of Harvard University. Research reported in this publication was supported by Office of the Director of the National Institutes of Health under award number DP5OD026389. The content is solely the responsibility of the authors and does not necessarily represent the official views of the National Institutes of Health.

